# Interactomics of SARS-CoV-2 Macrodomain 1 Reveals Putative Clients of ADP-ribosyl Hydrolase Activity

**DOI:** 10.1101/2025.10.21.683784

**Authors:** Crissey D. Cameron, Grace Heilmann, Lars Plate

## Abstract

Severe acute respiratory syndrome coronavirus 2 (SARS-CoV-2) has greatly impacted public health due to high rates of transmissibility and mutation during the COVID-19 pandemic. Macrodomain 1 (Mac1) of non-structural protein 3 remained well-conserved across variants and is critical for suppression of host immune response to infection, making Mac1 a promising target for therapeutic development. Mac1 binds and cleaves the post-translational modification ADP-ribose and is hypothesized to have a downstream effect on host interferon response, but the exact cellular targets of Mac1 are still unknown. Characterizing the substrates of Mac1 ADP-ribosyl hydrolase activity using a catalytically inactive mutant N40D can reveal critical virus-host interactions to identify protein targets of Mac1 and reveal mechanisms of host interferon suppression. Here, we co-immunoprecipitated WT Mac1 and Mac1 N40D from HEK293T and A549 cell lines and quantified changes in protein interactions by TMT-multiplexed tandem mass spectrometry. We identified interactions between Mac1 and ADP-ribosylated substrates involved in DNA damage response, cytoskeletal components, and cell cycle regulation. Additionally, several members of the TRiC complex involved in protein folding were selectively enriched with mutant Mac1 from A549 cells. These findings suggest a novel role of Mac1 in regulating host protein folding.

## Introduction

The COVID-19 pandemic, caused by severe acute respiratory syndrome coronavirus 2 (SARS-CoV-2), has resulted in a global health crisis and more than 7 million deaths.^1^ Transmission occurs primarily through respiratory droplets released by an infected individual. Once inhaled, the virus targets cells expressing angiotensin-converting enzyme 2 (ACE2), including epithelial and endothelial cells in the lungs.^2^ SARS-CoV-2 is a member of the *Coronaviridae* family and the *Betacoronavirus* genus, which also includes SARS-CoV, Middle East respiratory syndrome coronavirus (MERS-CoV), and murine hepatitis virus (MHV).^3^ Coronaviruses (CoV) are positive-sense, single-stranded RNA viruses that can be directly translated by host ribosomes.^3^ Following translation, viral nonstructural proteins (nsps) assemble into the replication-transcription complex (RTC) which drives genome replication within double membrane vesicles derived from virus-remodeled endoplasmic reticulum (ER).^4^

Two-thirds of the SARS-CoV-2 genome encodes for 16 nsps that are responsible for replication of the viral genome and disruption of key host cellular processes.^5^ Here, we investigate nsp3, the largest protein in the SARS-CoV-2 genome. Nsp3 contains 10-16 domains depending on the CoV variant, which are primarily responsible for ssRNA binding, ADP-ribose binding, and protease processing of the viral polypeptide.^6,7^ Nsp3 domains play a key role in overall coronavirus replication and infection.^7^ The papain-like protease (PLpro) in nsp3 which is responsible for cleaving the viral polyprotein and ISG15 modifications, has been the target for drug development, though none have been approved for treatment of SARS-CoV-2.^8–10^

Macrodomains (Mac) are highly conserved across many kingdoms of life and are found in a wide variety of species as well as positive sense RNA viruses.^11^ There are 3 macrodomains in nsp3: Mac 1, Mac 2 and Mac 3. Mac 1, located at position 207-386 in the N-terminal cytosolic region of nsp3, is primarily responsible for disrupting the interferon (IFN) response by binding to and hydrolyzing ADP-ribose (Figure 1A).^12,13^ All coronaviruses contain a conserved Mac1 domain and catalytic activity has been shown to be required for SARS-CoV-2 replication^6,14^. Mac2 and Mac3 are less conserved. They have RNA-binding activity, however, they are catalytically inactive and have not warranted further investigation for development of SARS-CoV-2 therapeutics.^15^ Given the essential role of Mac1 in immune evasion and high conservation across variants, this domain is a promising candidate for host-targeted therapeutic strategies aimed at disrupting conserved virus–host interactions.

**Figure 1:**
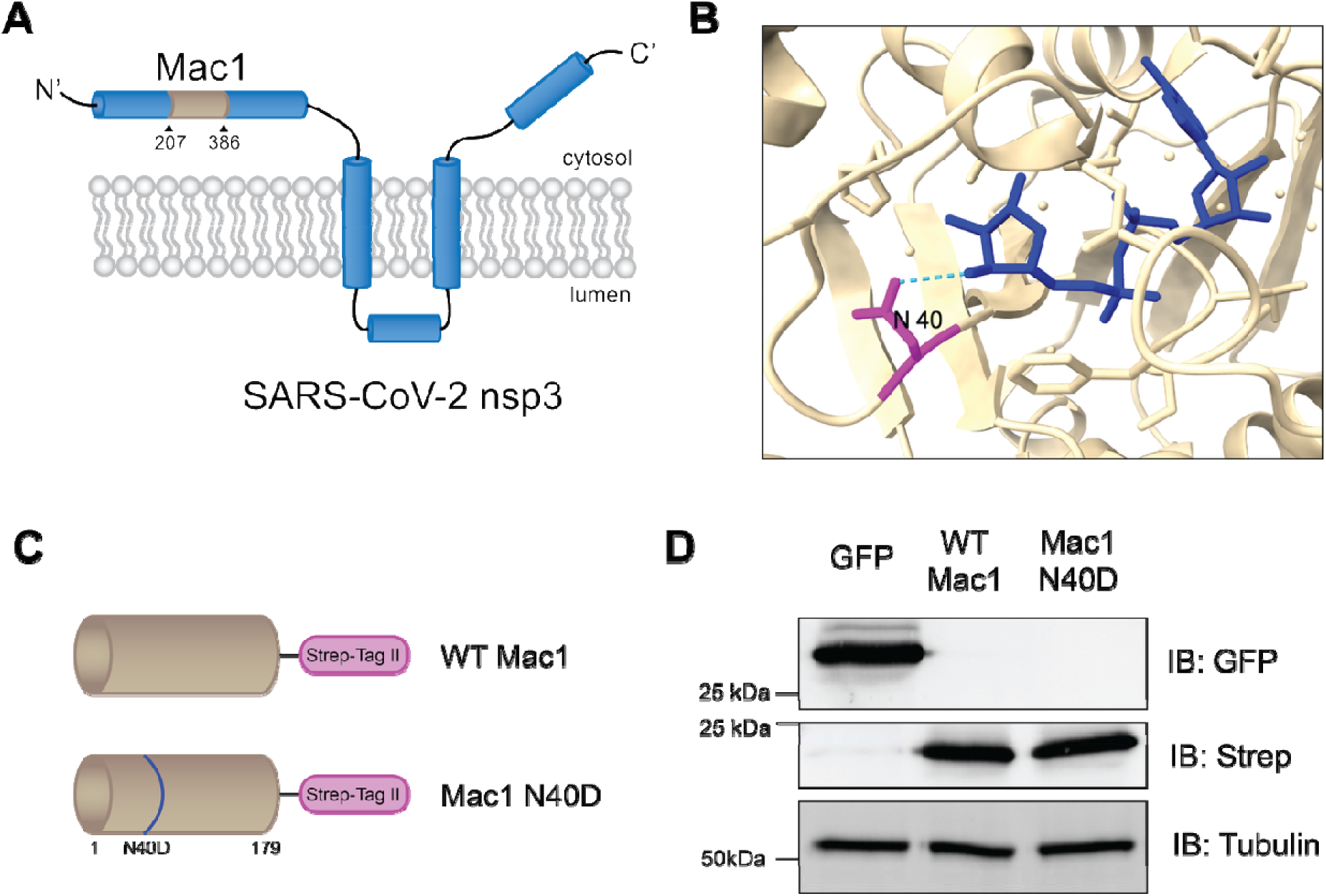
SARS-CoV-2 Mac1 and mutants can be used to study the role of ADP-ribosyl hydrolase activity *in cellulo*. A) The position of Mac1 in SARS-CoV-2 nsp3. B) Structure of monomer of Mac1 (tan) bound to ADP-ribose (dark blue, PBD:6WOJ).^12^ Asparagine (N) at position 40 (magenta) and hydrogen bond between N40 and ADP-ribose (light blue). C) Mac1 constructs used in this study. D) Western blot of HEK293T lysates in which Mac1 can be expressed as a single domain. Mac1 N40D expression is similar to WT Mac1.

ADP-ribosylation (ADPr) is a PTM introduced by poly-adenosine diphosphate-ribose polymerases (PARPs) and can either be one monomer: mono-ADP-ribose (MAR), or multiple ADP-ribose subunits: poly-ADP-ribose (PAR).^12,16,17^ ADPr modulates a significant number of downstream effects such as DNA repair, signal transduction, and control of epigenetics.^17^ It also plays an essential role in the host immune signaling pathway as several PARPs are activated by interferon signaling.^18,19^ Mac1 preferentially removes mono-ADP-ribose modification performed by PARP14 and the PARP9/DTX3L heterodimer which are activated in response to interferon stimulation.^12,20–22^ Mac1 is therefore critical for dampening the host immune response.^12^ This suggests that Mac1 plays an essential role in pathogenesis, specifically modulating the IFN response of the host cell. Inhibitors of Mac1 have been developed that directly bind the ADP-ribose site, though characterizing the substrates of Mac1 will open the doors for further development of host-targeted therapeutics.^23–25^

Mapping protein–protein interactions can provide critical insights into a protein’s function or substrate specificity by identifying its interaction partners within relevant cellular pathways. While no interactomics data have been collected for Mac1, interactors have been determined for nsp3.1, a truncation of nsp3 that contains the Mac1 domain. 411 high confidence interactors have been identified for nsp3.1, the majority of these being RNA-binding proteins.^26^ Other studies have investigated interactors of nsp3 variants, though none of the clinical variants have mutations in the Mac1 domain of nsp3.^27^

Hydrogen bonding between Mac1 N40 and the distal ribose moiety of an ADP-ribosylated substrate is necessary for catalytic activity of this domain (Figure 1B).^6,12,23,28,29^ Critically, Mac1 N40D retains ADP-ribose binding affinity despite having 100-fold reduction in catalytic activity, meaning that ADP-ribosylated clients of Mac1 will bind but will not be cleaved by the N40D mutant.^6^ By examining the differences in interactors between wildtype (WT) Mac1 and those with lengthened interactions with Mac1 N40D, we can identify host proteins that are potential ADP-ribosylated clients of this domain that preferentially interact with Mac1 N40D.

Here, we investigated the interactome of SARS-CoV-2 Mac1 and the catalytically inactive mutant Mac1 N40D to identify ADP-ribosylated substrates of this domain in two different cell types. In HEK293T cells, interactors were associated with cytoskeletal components, cell cycle regulation, and DNA damage response. Several proteins were identified that had greater interaction strength with Mac1 N40D when compared to WT Mac1 and may be potential substrates of ADP-ribosyl hydrolase activity: EIF4A1/2, NADK2, MEPCE, DTWD1, DPH2, and CSTF2T. In A549 lung carcinoma cells, a model cell line for SARS-CoV-2 infection, interactors were associated with viral IFN response and regulation of RNA splicing. With the addition of crosslinking in A549 cells, several members of the TRiC complex were identified as preferential interactors of Mac1 N40D, suggesting that Mac1 has ADP-ribosyl hydrolase activity against members of the complex, possibly with a regulatory role in host protein folding.

## Results

### Macrodomain 1 and mutants can be expressed and purified from cell lysate

To investigate the key interactors of the Mac1 domain of SARS-CoV-2, we generated a plasmid expressing WT Mac1 that had been truncated from the full-length nsp3 and contained a *C*-terminal Strep-Tag II (ST). An N40D mutant was also generated to serve as a catalytically inactive control that will recognize and retain substrates (Figure 1C). The two constructs were transiently transfected in human embryonic kidney (HEK293T) cells and exhibited similar levels of expression (Figure 1D).^6^

### Interactors of Mac1 were determined in HEK293T cells

Interactors of Mac1 were first determined in HEK293T cells. Mac1 and interactors and binding partners were enriched from lysates expressing GFP, WT Mac1, or Mac1 N40D using the *C*-terminal Strep-Tag (Figure 2A), and the isolation was confirmed by Western blot (Figure S1A). Enriched samples were prepared for bottom-up liquid chromatography - tandem mass spectrometry (LC-MS/MS) analysis and were labeled with tandem mass tag (TMTpro 16-plex) reagents which allowed for multiplexing and quantitative comparison. Following MS analysis, more than 2,000 proteins were identified, and 74 proteins passed cutoffs of enrichment greater than 2 standard deviations (SD) and significance greater than p=0.05 when comparing abundance of proteins in the Mac1 samples to a GFP negative control (Figures S1B-C, Table 1). The 74 enriched proteins, hereafter called interactors, were significantly associated with the cytoskeleton, cytoplasm, extracellular region, and the mitochondrion when analyzed for subcellular localization with SubcellulaRVis (Table 1).^30^ Interactors were also analyzed for biological process gene ontology (GO) term enrichment using EnrichR^31^ and were found to be associated with cytoskeleton organization and transport, and DNA repair (Figure 2B, Table 1). Interactors were also associated with guanosine triphosphate (GTP) and ADP binding biological processes GO terms (Figure 2C, Table 1). When comparing the interactors identified in this study to previously identified interactors of nsp3.1, a truncated analog of nsp3 containing the Mac1 domain, there was an overlap of three proteins: TUBB4B, SCL25A5, and POLD3 (Figure S1D).^26^ Interactors were also compared against a dataset of proteins with ADP-ribose modification under stress and there was an overlap of eight proteins, including DTL, EIF4A2, and MEPCE (Figure S1D).^32^

**Figure 2:**
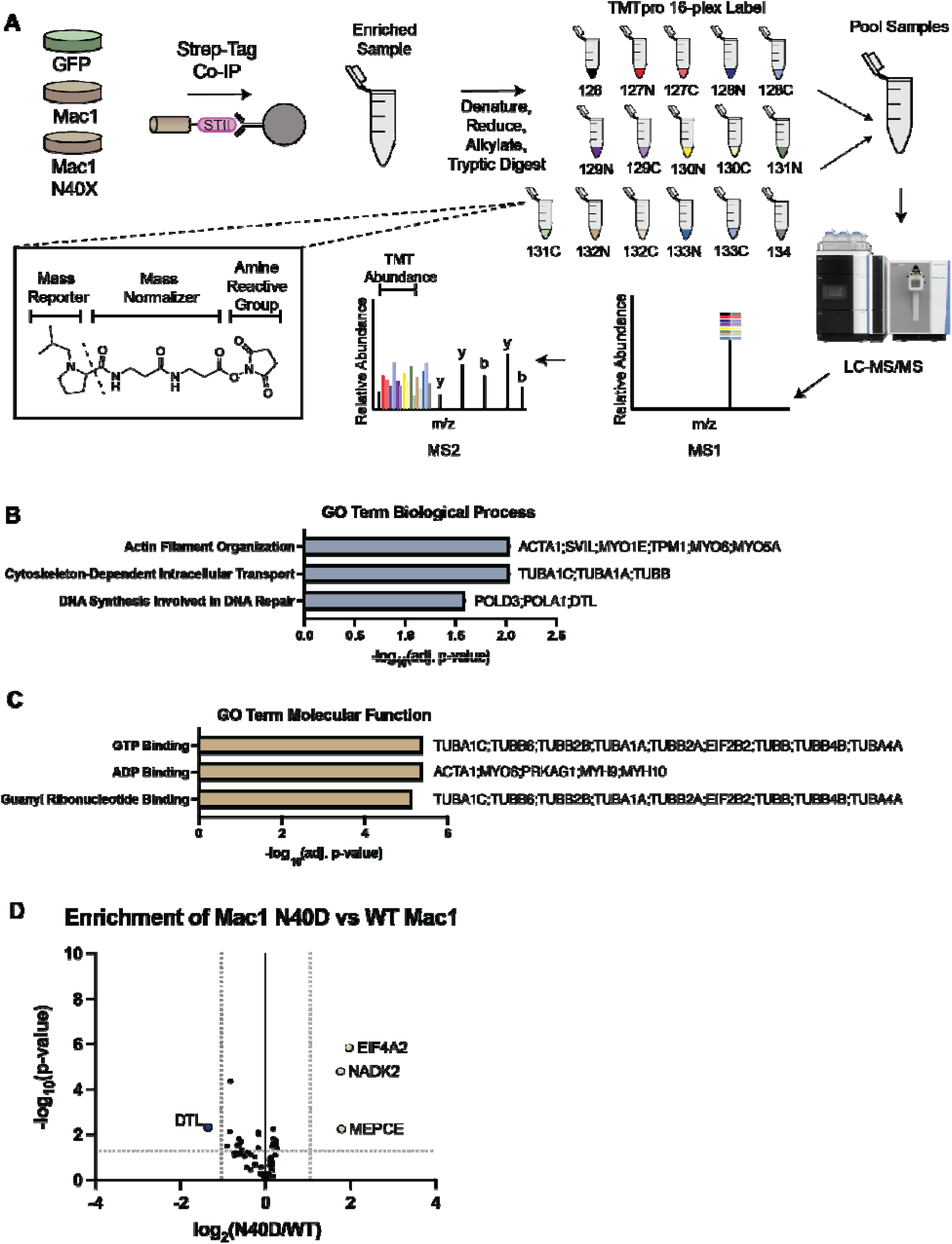
Mac1 is associated with cytoskeletal and DNA repair proteins in HEK293T cells. A) Workflow schematic for enrichment and LC-MS/MS analysis of Mac1 and mutants expressed in HEK293T cells. Cells expressing GFP, WT Mac1 or Mac1 N40D were lysed and subjected to co-immunoprecipitation using Strep-tactin resin, which binds the 2x Strep-Tag on the *C*-terminus. Enriched Mac1 and interactors were then reduced, alkylated, and digested using Trypsin/LysC and then labeled with TMTpro 16-plex reagents for later deconvolution of MS quantification data. Samples were pooled and analyzed using liquid chromatography-tandem mass spectrometry to identify and quantify peptides. B) Biological process gene ontology analysis for 74 combined interactors of WT Mac1 and Mac1 N40D in HEK293T cells. Interactors are associated with cytoskeletal organization and DNA repair. C) Molecular function gene ontology analysis for 74 combined interactors of WT Mac1 and Mac1 N40D in HEK293T cells. Interactors have GTP and ADP binding activity. D) Volcano plot comparing the enrichment of all 74 interactors between WT Mac1 and Mac1 N40D. EIF4A2, NADK2, and MEPCE were more enriched with Mac1 N40D, and DTL was more enriched with WT Mac1.

We next wanted to assess if there was any impact of the N40D mutation on the interactors of Mac1. We compared the abundances of the interactors identified in this study between WT Mac1 and Mac1 N40D and found that four proteins were differentially enriched: DTL was more enriched with WT Mac1, and MEPCE, EIF4A2, and NADK2 were more enriched with Mac1 N40D (Figure 2D, S1E).

### Covalent Capture of Mac1 Interactors Reveals Association with Cell Cycle Regulation and DNA Repair Machinery

Interactions of host substrates with Mac1 are expected to be short-lived and transient in nature. To capture these transient interactions and observe them by MS, we used a covalent crosslinker, dithiobis(succinimidyl propionate) (DSP), to covalently link nearby primary amines in intact cells prior to lysis. Proteins that interact with Mac1 will be covalently bound to Mac1 and should co-immunoprecipitate during enrichment. The same procedure as above was performed with GFP, WT Mac1, and Mac1 N40D in HEK293T cells with the addition of DSP crosslinker to capture transient interactions. The enrichment was confirmed by Western blot (Figure S2A), and proteins were identified and quantified by LC-MS/MS.

More than 3,200 proteins were identified in this dataset with greater than two peptides per protein and 72 proteins passed enrichment cutoffs for interactors (Figure S2B,C, Table 2). The interactors from this crosslinked dataset were associated with the cytoskeleton, cytoplasm, and nucleus components when searched with SubcellulaRVis (Table 2).^30^ The interactors were also analyzed for gene ontology (GO) term enrichment using EnrichR, and associations with DNA repair and cell cycle transition were uncovered, two processes known to be influenced by ADP-ribosylation (Table 2).^31^

Due to crosslinking of nearby proteins during the sample preparation, enrichment of Mac1 and interactors may result in co-immunoprecipitation of large complexes that have been covalently linked together. Visualizing this in the form of a network plot can be useful for understanding the known interaction hierarchies identified within the set of interactors (Figure 3A). Here, we identified 23 interactors associated with regulation of the cell cycle and 7 interactors involved in regulation of DNA damage response. Many of the interactors involved in the cell cycle are known to be associated and are connected on the network plot. In addition, three interactors involved in regulation of DNA damage response cluster together: SMARCD2, ARID1A, and BCL7A. These proteins are subunits of the SWI/SNF complex that is involved in chromatin remodeling and regulation of gene expression.^33^

**Figure 3:**
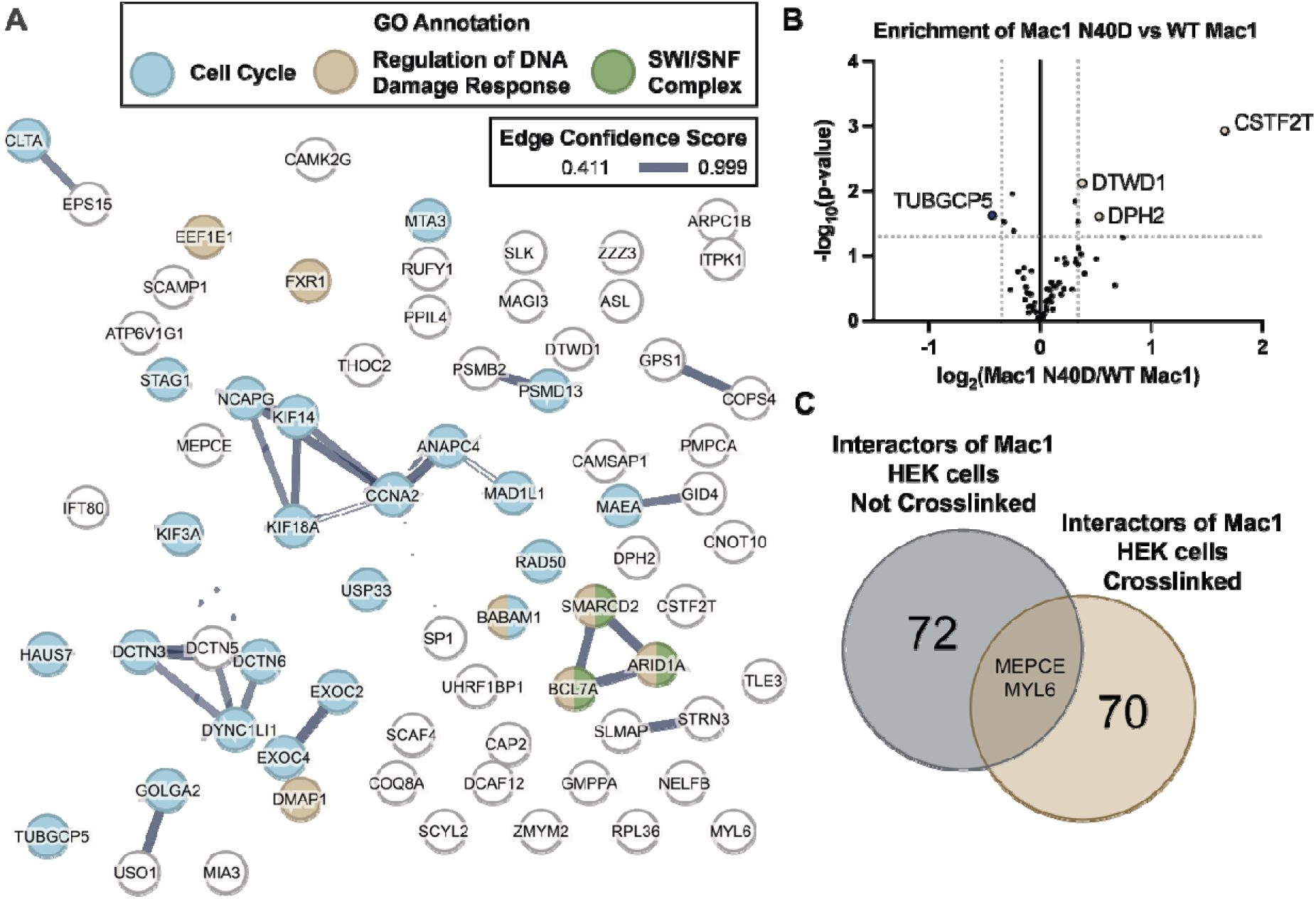
Crosslinked interactors of SARS-CoV-2 Mac1 in HEK293T cells are associated with cell cycle regulation and DNA damage response. A) Network analysis of interactors from crosslinked HEK293T interactomics of SARS-CoV-2 Mac1. Interactors associated with cell cycle (GO:0007049, 23/72) highlighted in teal and interactors associated with positive regulation of response to DNA damage stimulus (GO:2001022, 7/72) highlighted in tan. Edge thickness represents the confidence score that the two nodes are associated from experimental data and database annotation by STRING analysis. B) Volcano plot comparing the relative enrichment of interactors between Mac1 N40D and WT Mac1. Interactors with greater enrichment with Mac1 N40D highlighted in brown and interactor with greater enrichment with WT Mac1 highlighted in blue. C) Venn diagram showing the overlap between interactors from crosslinked (brown) and not crosslinked (grey) interactomics from HEK293T cells in this study.

We next wanted to understand the impact of the N40D mutation on Mac1 interactions. The 72 interactors identified from both WT and mutant Mac1 were analyzed for differential enrichment (Figure 3B). Three proteins were more highly enriched with Mac1 N40D: DTWD1, DPH2, and CSTF2T. One protein was more highly enriched with WT Mac1: TUBGCP5. Two interactors were identified as enriched in both this crosslinked dataset and the dataset acquired without crosslinking presented in Figure 2: MEPCE and MYL6 (Figure 3C).

### Mac1 in A549 lung cells interacts with TRiC complex

To obtain more biologically relevant data regarding SARS-CoV-2 nsp interactions in lung cells, we performed enrichment of Mac1 with and without DSP crosslinker in A549 cells, a human lung carcinoma cell line commonly used to study this respiratory virus. A549 cells were transiently transfected with GFP, WT Mac1 or Mac1 N40D. Lysates were enriched for the Strep-Tag on Mac1 and analyzed by multiplexed LC-MS/MS.

In samples without crosslinking (Figure S3A), 549 proteins were identified (Table 3). When comparing Mac1-containing samples to the GFP negative control, Mac1 was highly enriched for both the WT and mutant (Figure S3B-C). Only three proteins for each Mac1 passed interactor cutoffs of 2 SD enrichment and significant p-value: HNRNPH2, EIF4A1, and YTHDF3 for WT and HSPA1A, ESYT2, and SAMHD1 for Mac1 N40D. Due to the low number of proteins identified and the low number of interactors, a less stringent cutoff of 1 SD enrichment was used, hereafter termed “enriched proteins”. These enriched proteins from the un-crosslinked data revealed associations of Mac1 with viral infection pathways, interferon and immune signaling (Figure 4A, Table 3).^31^

**Figure 4:**
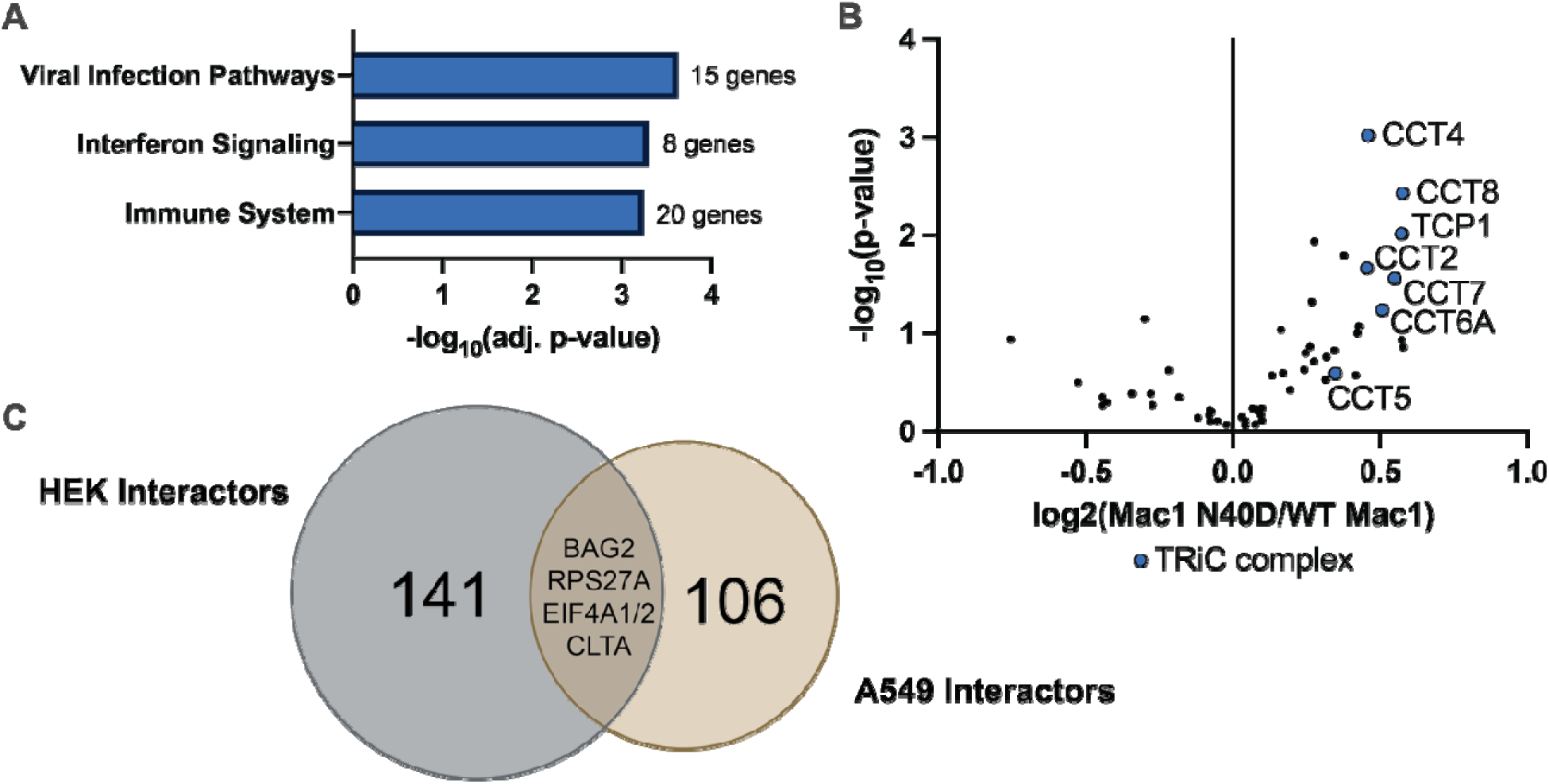
Enriched proteins with Mac1 from A549 cells are associated with viral infection and the TRiC complex. A) Reactome GO Term analysis from Mac1 enrichment without crosslinking from A549 cell lysate. Proteins with greater than 1 SD enrichment are associated with viral infection pathways, interferon signaling, and immune signaling. B) Volcano plot of proteins with greater than 1 SD enrichment from crosslinked Mac1 enrichment compared to GFP plotted comparing Mac1 N40D and WT Mac1 to determine differential enrichment between WT and mutant. Of 55 proteins that were enriched with WT Mac1 and/or Mac1 N40D, 7 are components of the TRiC complex. In differential enrichment analysis, all 7 of these TRiC proteins were more enriched with Mac1 N40D when compared to WT Mac1. C) Venn diagram comparing the interactors of Mac1 identified in HEK293T cells to the enriched proteins from A549 cells, crosslinked and un-crosslinked combined. Four proteins were enriched in both datasets: BAG2, RPS27A, EIF4A1/2, and CLTA.

When DSP crosslinker was used to capture transient interactions with Mac1, 435 proteins were identified from Mac1-enriched samples from A549 cells (Figure S3D, Table 4). Comparisons between Mac1 samples and the GFP negative enrichment control showed the Mac1 bait was the most enriched protein for both WT and N40D samples (Figure S3E-F). Again, only three proteins passed interactor cutoffs of 2 SD enrichment and significant p-value: MYL12B and SPTBN1 for both WT and N40D, and CLTC which was unique to WT Mac1 (Figure S3E-F). All three interactors have been observed with ADPr modification in previous studies.^34–38^ Enriched proteins were determined using 1 SD cutoff, and those enriched proteins were analyzed for differential enrichment between WT and N40D (Figure 4B). Of the 61 proteins that were enriched with Mac1, seven are members of the T-complex protein Ring Complex (TRiC): CCT2, CCT4, CCT5, CCT6A, CCT7, CCT8, and TCP1. All enriched members of the TRiC complex are more enriched with Mac1 N40D than WT Mac1. All enriched components of the TRiC complex have also been shown to have ADPr modification, though the effects of ADPr on TRiC function has not been fully explored.^34–36,39–43^

The amount of overlap between crosslinked and un-crosslinked enriched proteins in A549 cells was relatively low. Five proteins were identified as enriched both with and without crosslinking: CCT2, CLTC, EPRS, FLNA, and VCP (Figure S3G). Through searching the ADPriboDB 2.0^34^, four of these proteins were found to have ADPr modification: CCT2, CLTC, FLNA, and VCP.^36,38,39,42,43^ The overlap between enriched proteins from A549 cells and interactors identified in HEK293T cells was also relatively low. Importantly, four proteins were identified as co-enriched with Mac1 in both cell types: BAG2, RPS27A, EIF4A1/2, and CLTA, all of which are known to have ADPr modification.^34–36,39,40,42–44^ These conserved interactions across cell types could possibly point to ADP-ribosylated clients of Mac1.

## Discussion

This work characterizes the interactome of the SARS-CoV-2 nsp3 Mac1 domain in HEK293T cells and A549 cells with and without covalent capture. Without the use of crosslinker, Mac1 interactors in HEK293T cells were associated with cytoskeletal components and displayed GTP and ADP binding. ADP-ribosylation by PARP9 and PARP14 are known to make critical ADPr modifications to cytoskeletal components that are responsible for cell shape and cell movement.^45^ Mac1 having ADP-ribosyl hydrolase activity against modifications made by PARPs 9 and 14 suggests that the interactions identified here between Mac1 and cytoskeletal components may be Mac1 substrates.

When interactors were compared for differential enrichment, DTL was more enriched with WT Mac1 and MEPCE, EIF4A2, and NADK2 were more enriched with Mac1 N40D. The WT Mac1-specific interactor denticleless E3 ubiquitin protein ligase homolog (DTL) is involved in cell cycle regulation, response to DNA damage, and protein ubiquitination.^46^ In a recent study using transcriptomics data from SARS-CoV-2 infected patients, DTL was shown to be upregulated in the viral condition, and was identified as a hub protein that has the potential to be used as a biomarker for infection.^47^ Overexpression of DTL has been shown to increase host genomic instability by interfering with the non-homologous end-joining (NHEJ) repair mechanism in response to double-stranded breaks (DSB).^46^ DTL has also been shown to be ADP-ribosylated.^34,39,48^

The Mac1 N40D-specific interactors MEPCE, EIF4A2, and NADK2 are associated with DNA repair, translation, and mitochondrial function respectively. MEPCE, or methylphosphate capping enzyme, is involved in double-stranded break (DSB) repair.^49^ While some evidence exists that MEPCE is itself ADP-ribosylated,^35,39,48^ DSB repair is well known to be regulated by ADPr through recruitment of repair factors and chromatin remodelers.^50^ Mac1 interaction with MEPCE may suggest a role in regulation of DNA damage that can be caused by SARS-CoV-2 infection.^51^ EIF4A2 is an RNA helicase that is part of the eIF4F translation initiation complex and is known to be upregulated during SARS-CoV-2 infection in response to stress.^52^ One study identified EIF4A2 as ADP-ribosylated using 6-alkyne adenosine labeling in MDA-MB-231 breast cancer cells.^35^ NADK2 is a mitochondrial enzyme that is responsible for maintaining supply of NADPH essential for mitochondrial function.^53^ It is not known to be ADP-ribosylated or have any role in viral infection. Mac1 is also not known to localize to the mitochondria. These data together suggests that NADK2 interaction with Mac1 may not be meaningful.

Interactors in the HEK293T dataset were highly associated with cell cycle regulation and DNA damage response when crosslinking was conducted, indicating that Mac1 has transient interactions with these machineries. In the HEK293T crosslinked dataset, TUBGCP5 protein was enriched more with WT Mac1 than Mac1 N40D. TUBGCP5 or gamma-tubulin complex component 5 is part of the gamma tubulin complex responsible for nucleation at the centrosome during cell division.^54^ Interactions between SARS-CoV-2 proteins M and ORF3a and members of the gamma tubulin complex have been characterized and are thought to lengthen the cell cycle arrest and apoptosis.^55^ While TUBGCP5 is not known to be ADP-ribosylated, cytoskeletal and cell cycle regulation proteins commonly have this PTM. The Mac1 N40D-specific interactors from the crosslinked HEK293T dataset, DTWD1, DPH2, and CSTF2T, are involved in tRNA stability, protein elongation, and pre-mRNA processing.^56^ Though none of these three proteins are known to be modified with ADP-ribose, each are involved in critical cellular processes that are often associated with ADPr.^34^

In A549 lung carcinoma cells, proteins co-enriched with Mac1 without DSP crosslinking were associated with viral infection and immune system activation. Notably, the ADP-ribosylated substrate of DPH2 was identified, EEF2, as well as other proteins involved in translation: EIF4A1, EIF2AK2, and RPS. Isoform EIF4A2 was identified as an interactor of Mac1 in HEK293T cells, suggesting conserved engagement of this host RNA helicase that is part of the pre-initiation complex. Proteins involved in immune signaling were also identified such as YWHAZ, SAMHD1, and TBL1XR1, the first two of which are ADP-ribosylated.^42,43^

Notably in the set of proteins co-enriched with Mac1 from the crosslinked A549 cells, seven members of the TRiC complex were identified, and all the observed TRiC proteins were more enriched with Mac1 N40D than WT Mac1. TRiC is responsible for folding approximately 10% of the proteome, particularly cytoskeletal components including actin and tubulin.^57–59^ Though interactions between nsp3 and TRiC have not been previously identified, TRiC has been shown to interact with other SARS-CoV-2 proteins, namely nsp12, and to be critical for viral fitness during infection.^60^ TRiC has also been shown to be responsible for folding reovirus capsid proteins which is required for viral particle assembly.^61^ Since Mac1 N40D has expression and stability similar to WT Mac1^6^, it is unlikely that the increased interactions of Mac1 N40D with TRiC indicate a substrate folding relationship. Rather, increased interactions of Mac1 N40D with TRiC components indicate a regulatory role of TRiC ADP-ribosylation. Investigating the ADPr state of CCT proteins could reveal a key regulatory mechanism of protein folding in response to cellular stress.

Our interactomics of Mac1 of SARS-CoV-2 in HEK293T and A549 cells has identified possible substrates for Mac1 ADP-ribosyl hydrolase activity. Following validation of these interactions, functional analysis via protein knockdown should be performed to assess the impact of these interactions on immune response and viral fitness. The interactors identified here implicate Mac1 in cytoskeletal organization, DNA damage response, and protein folding via the TRiC complex.

## Acknowledgements

This work was supported by the National Institute of General Medical Sciences (2R35GM133552, **Lars Plate,** 5T32GM149371, **Crissey Cameron**), the National Science Foundation’s Mathematical and Physical Sciences Program (2349507, **Grace Heilmann**), and Vanderbilt University startup funds.

## List of Tables

Table 1: Proteins, SubcellulaRVis, and GO Term analysis from HEK293T Mac1 Interactomics without Crosslinker

Table 2: Proteins, SubcellulaRVis and GO Term analysis from HEK293T Mac1 Interactomics with Crosslinker

Table 3: Proteins and GO Term analysis from A549 Mac1 Interactomics without Crosslinker

Table 4: Proteins from A549 Mac1 Interactomics with Crosslinker

Table 5: Primers used for site-directed mutagenesis of SARS-CoV-2 Mac1

## Methods

### Site Directed Mutagenesis

Using the nsp3.1 truncated sequence of SARS-CoV-2 isolate Wuhan-Hu-1 MN908947 nsp3 generated in Almasy et al.^26^, the plasmid was further truncated to isolate macrodomain 1 (Mac1) using primers in Table 5 using Q5 site-directed mutagenesis according to manufacturer protocols (NEB, E0554S). Site directed mutagenesis was then conducted using PFU Ultra II Fusion HS DNA polymerase (Agilent, 600670) according to the manufacturer’s protocol to introduce N40D mutation. Plasmids were purified using Plasmid Plus Midi Kit (Qiagen, 12943) from DH5α competent *E. coli*.

### Cell Culture and Transfection

HEK293T and A549 cells were grown using Dulbeccos’ Modified Eagle’s Medium (high glucose) supplemented with 10% fetal bovine serum, 1% penicillin/streptomycin and 1% glutamine. Cell cultures were maintained at 37°C and 5% CO_2_. HEK293T cells were transiently transfected using calcium phosphate method with wildtype Mac 1 (WT), mutant Mac1 (N40A or N40D) or green fluorescent protein (GFP). Media was changed 16 hours post-transfection and the cells were harvested after 24 hours. A549 cells were transiently transfected with using FuGENE 4k (FuGENE, 4K-1000) according to manufacturer’s protocol with GFP, WT Mac1, or Mac1 N40D plasmid in antibiotic-free medium. Media was replaced with antibiotic-supplemented media 16 hours post transfection and the samples were harvested after 24 hours.

### Harvesting

Cells were first washed with phosphate buffered saline (PBS) and then scraped and collected in cold 1 mM ethylenediaminetetraacetic acid (EDTA) in PBS. Cells were then pelleted at 200 x g for 5 minutes at 4°C and washed twice with PBS. The remaining PBS was removed by aspiration. Cells were lysed in 200 µL TNI buffer (50 mM Tris pH 7.5, 150 mM NaCl, 0.5% IGEPAL-CA-630) supplemented with Roche cOmplete protease inhibitor on ice for 15 minutes followed by 15 minutes sonication at room temperature. Lysates were cleared by centrifugation at 21.1 x g for 15 minutes at 4°C. Protein concentration was determined using BCA assay (Thermo, 23225) and normalized to the lowest concentration in the sample set.

### DSP Crosslinking

For crosslinking experiments, cells were washed with PBS, scraped and collected in a tube, and washed with PBS again as described above. DSP (Fisher, PI22585) was dissolved in DMSO for a 50 mM 100x stock immediately before use. DSP was diluted 100x in 1 mL PBS for a working concentration of 0.5 mM, added directly to cells, and incubated rocking for 30 minutes at room temperature. The DSP was then quenched with 100 µL of 1 M Tris pH 7.5 and incubated for 15 minutes at room temperature. Two washes were performed with PBS and lysis procedure was performed as described previously with TNI lysis buffer.

### Co-Immunoprecipitation

4B Sepharose beads (Sigma-Aldrich, 4B200) and Streptactin beads (IBA, 2-5030-002) were pre-washed 4x with TNI buffer. Lysates were pre-cleared with 20 µL 4B Sepharose beads slurry per sample rotating for 1 hour at 4°C. Beads were cleared at 400 x g for 5 min at 4°C and pre-cleared lysate was added to 20 µL of FLAG bead slurry per sample, rotating overnight at 4°C. Resin was washed 4x with TNI buffer, cleared by centrifugation at 400 x g, and dried with 30G needle after the final wash. Resin-bound proteins were eluted twice in 3x modified Laemelli buffer (6% SDS, 62.5 mM Tris pH 6.8) for 30 minutes at room temperature and 15 minutes at 37°C.

### Western Blot

Normalized lysates were reduced with 6x SDS Laemelli buffer (12% SDS, 125 mM Tris pH 6.8, 20% glycerol, bromophenol blue, 100 mM DTT) and heated to 95°C for 5 minutes before being run on 12% SDS-PAGE gel at 200 V for 1 hour. The gels were then transferred to PVDF membrane (Millipore, IPFL00010) using wet transfer at 100 V for 80 minutes. Blots were probed with 1:1000 dilutions in 5% bovine serum albumin (RPI, A30075100), 0.1% sodium azide in PBS of anti-GFP (Vanderbilt Antibody and Protein Resource Core clone #1C9A5), anti-StrepII [FITC] mAb mouse (Genscript, A01736), and hFAB Rhodamine anti-Tubulin (Bio-Rad, 12004165). All Western blot images were acquired on BioRad ChemiDoc MP using autorapid settings.

### Mass Spectrometry Sample Preparation

Eluted proteins were precipitated via methanol/chloroform water (3:1:3) and washed 3x with methanol. Pellets were air-dried before resuspension with Rapigest SF (Waters) followed by the addition of 10 µL of 0.5 HEPES (pH 8.0). Volume was then adjusted to 47.5 µL with H2O. Proteins were reduced with 5 mM tris(2-carboxyethyl)phosphine (TCEP) (Sigma, 75259) for 30 minutes at room temperature followed by alkylation with 10 mM iodoacetamide (Sigma, I6125). Proteins were digested overnight at 37°C shaking at 750 rpm using 0.5 µg of Trypsin/Lys-C protease mix (Pierce, A40007). TMTpro 16plex reagents (Thermo Fisher, 44520) in 40% v/v acetonitrile were used to label the digested peptides for 1 hour at room temperature and quenched with 0.4% w/v ammonium bicarbonate for 1 hour at room temperature. Samples were pooled acidified using 5% v/v formic acid (Fisher, A117) to reach a pH of 2. The sample was then concentrated to 1/3 of its volume using a speedvac and resuspended in Buffer A (97% water, 2.9% acetonitrile, 0.1% formic acid, v/v/v). Cleaved Rapigest SF was removed by centrifugation for 30 minutes at 21,000 x g.

### Liquid Chromatography-Mass Spectrometry

Multidimensional Protein Identification Technology (MudPIT) microcolumns were prepared as previously described.^62^ Peptide samples were directly loaded onto the columns using a high-pressure chamber. Samples were then desalted for 30 minutes with buffer A (97% water, 2.9% acetonitrile, 0.1% formic acid v/v/v). LC-MS/MS analysis was performed using an Exploris480 (Thermo Fisher) mass spectrometer equipped with an Ultimate3000 RSLCnano system (Thermo Fisher). MudPIT experiments were performed with 10 μL sequential injections of 0, 10, 30, 60, and 100% buffer C (500 mM ammonium acetate in buffer A), followed by a final injection of 90% buffer C with 10% buffer B (99.9% acetonitrile, 0.1% formic acid v/v) and each step followed by a 92 minute gradient from 5% to 35% B and a short column flush up to 85% B for 7 minutes with a flow rate of 500 nL/minute on a 20 cm fused silica microcapillary column (ID 100 μm) ending with a laser-pulled tip filled with Aqua C18, 3 μm, 125 Å resin (Phenomenex). Electrospray ionization (ESI) was performed directly from the analytical column by applying a voltage of 2.2 kV with an inlet capillary temperature of 275°C. Data-dependent acquisition of mass spectra was carried out by performing a full scan from 400-1600m/z at a resolution of 120,000. Top-speed data acquisition was used for acquiring MS/MS spectra using a cycle time of 3 seconds, with a normalized collision energy of 36, 0.4 m/z isolation window, automatic maximum injection time and 100% normalized AGC target, at a resolution of 45,000 and a defined first mass (m/z) starting at 110.

Peptide identification and TMT-based protein quantification was carried out using Proteome Discoverer 2.4. MS/MS spectra were extracted from Thermo Xcalibur .raw file format and searched using SEQUEST against a Uniprot human proteome database (accessed 03/2014 and containing 28,860 entries) supplemented with the SARS-CoV-2 Macrodomain 1 sequence. The database was curated to remove redundant protein and splice-isoforms. Searches were carried out using following parameters: 20 ppm peptide precursor tolerance, 0.02 Da fragment mass tolerance, minimum peptide length of 6 amino acids, trypsin cleavage with a maximum of two missed cleavages, dynamic methionine modification of +15.995 Da (oxidation), dynamic protein N-terminus +42.011 Da (acetylation), −131.040 (methionine loss), −89.030 (methionine loss + acetylation), static cysteine modification of +57.0215 Da (carbamidomethylation), and static peptide N-terminal and lysine modifications of +304.2071 Da (TMTpro 16plex).

The mass spectrometry proteomics data have been deposited to the ProteomeXchange Consortium via the PRIDE^63^ partner repository with the dataset identifier PXD069398 and 10.6019/PXD069398.

### Mass Spectrometry Data Analysis

Raw TMT abundances from Proteome Discoverer were median normalized and log_2_ transformed within each 16-plex. The expression changes between Mac1 samples and DMSO samples were calculated for each protein and tested for significance with a paired t-test. Proteins with a log_2_(fold change) greater than two standard deviations away from 0 and p-value less than 0.05 were considered interactors and were combined from WT Mac1 and Mac1 N40D comparisons.

