## Supplemental Information for "Interactomics of SARS-CoV-2 Macrodomain 1 Reveals Putative Clients of ADP-ribosyl Hydrolase Activity"

**for**

### Supplemental Information

**Figure S1:** Mac1 and interactors can be successfully enriched, and the identified proteins overlap with previously identified interactors. 3
**Figure S2:** Crosslinked Mac1 in HEK293T cells can be enriched using Co-IP. 5
**Figure S3:** Mac1 enrichment from A549 cells. 7

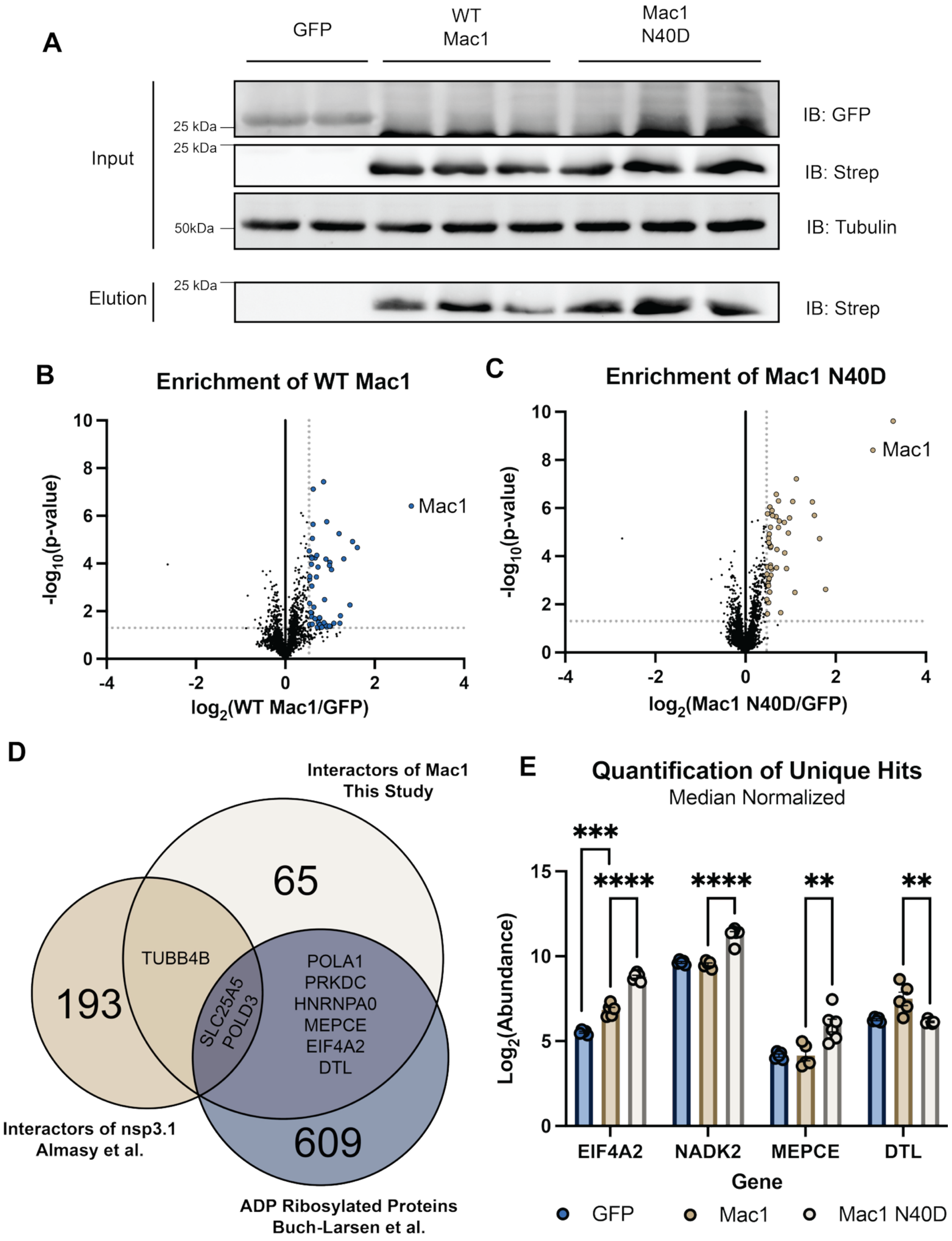
A
GFP
WT Mac1
Mac1 N40D
Input
Elution
25 kDa
25 kDa
50kDa
25 kDa
IB: GFP
IB: Strep
IB: Tubulin
IB: Strep
B
Enrichment of WT Mac1
Mac1
-log10(p-value)
log2(WT Mac1/GFP)
C
Enrichment of Mac1 N40D
Mac1
-log10(p-value)
log2(Mac1 N40D/GFP)
D
Interactors of Mac1
This Study
65
193
TUBB4B
SLC25A5
POLD3
POLA1
PRKDC
HNRNPA0
MEPCE
EIF4A2
DTL
609
Interactors of nsp3.1
Almasy et al.
ADP Ribosylated Proteins
Buch-Larsen et al.
E
Quantification of Unique Hits
Median Normalized
***
****
****
**
**
Log2(Abundance)
EIF4A2
NADK2
MEPCE
DTL
Gene
GFP
Mac1
Mac1 N40D

**Figure S1:** Mac1 and interactors can be successfully enriched, and the identified proteins overlap with previously identified interactors. A) Western blot analysis of Strep-Tag enrichment from HEK293T cells expressing GFP, WT Mac1 or Mac1 N40D. Prior to enrichment (input), Western blot analysis shows that cells express the expected protein of interest (GFP or Mac1), as demonstrated by anti-GFP and anti-Strep-Tag blots. Following enrichment and elution from the Strep-tactin beads (elution), Mac1 and the mutant were successfully enriched, as shown by the anti-Strep signal. B) Volcano plot comparing enrichment of identified proteins between WT Mac1 and GFP negative enrichment control. The Mac1 bait is the most enriched protein. C) Volcano plot comparing enrichment of proteins between Mac1 N40D and GFP negative enrichment control. The Mac1 bait is the second-most enriched protein following EIF4A2. D) Venn diagram comparing the interactors from this study (light tan) to interactors of the truncated version of nsp3 that contains the Mac1 domain, nsp3.1, from Almasy et al.[26] (tan), and proteins known to have ADP-ribose modification from Buch-Larsen et al.[32] (blue). E) Quantification from n = 5-6 of the $\log_2$(Abundance) of interactors that selectively interact with WT Mac1 (DTL) or Mac1 N40D (EIF4A2, MEPCE, NADK2). Multiple unpaired t-test performed with $\alpha = 0.05$.

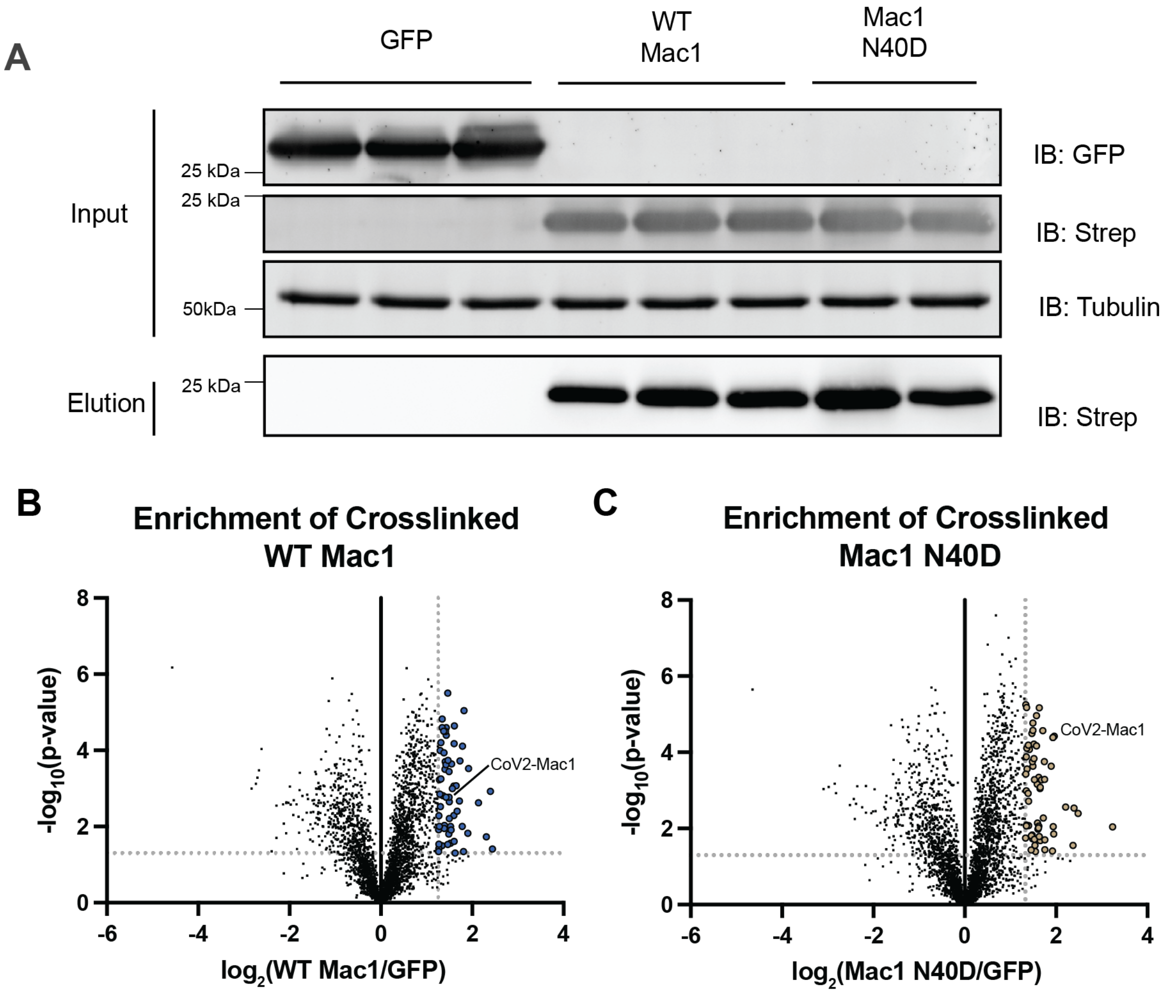


**Figure S2:** Crosslinked Mac1 in HEK293T cells can be enriched using Co-IP. A) Representative Western blot showing signal from GFP, Strep-Tag (represents Mac1), and tubulin from samples before (input) and after (elution) enrichment for Strep-Tag. HEK293T cells were transiently transfected with GFP, WT Mac1 or Mac1 N40D and crosslinked using DSP. Co-immunoprecipitation was then performed to enrich Strep-Tag and crosslinked interactors. Strep signal in the elution sample in the WT Mac1 and Mac1 N40D lanes indicates that the enrichment was successful. B) Volcano plot comparing the relative abundance of all identified proteins between WT Mac1 and GFP samples. Proteins that display greater than 2 standard deviation enrichment and greater than p=0.05 confidence in the WT Mac1 sample are interactors highlighted in blue. C) Volcano plot comparing the relative abundance of all identified proteins between Mac1 N40D and GFP samples. Proteins that display greater than 2 standard deviation enrichment and greater than p=0.05 confidence in the Mac1 N40D sample are interactors highlighted in brown.

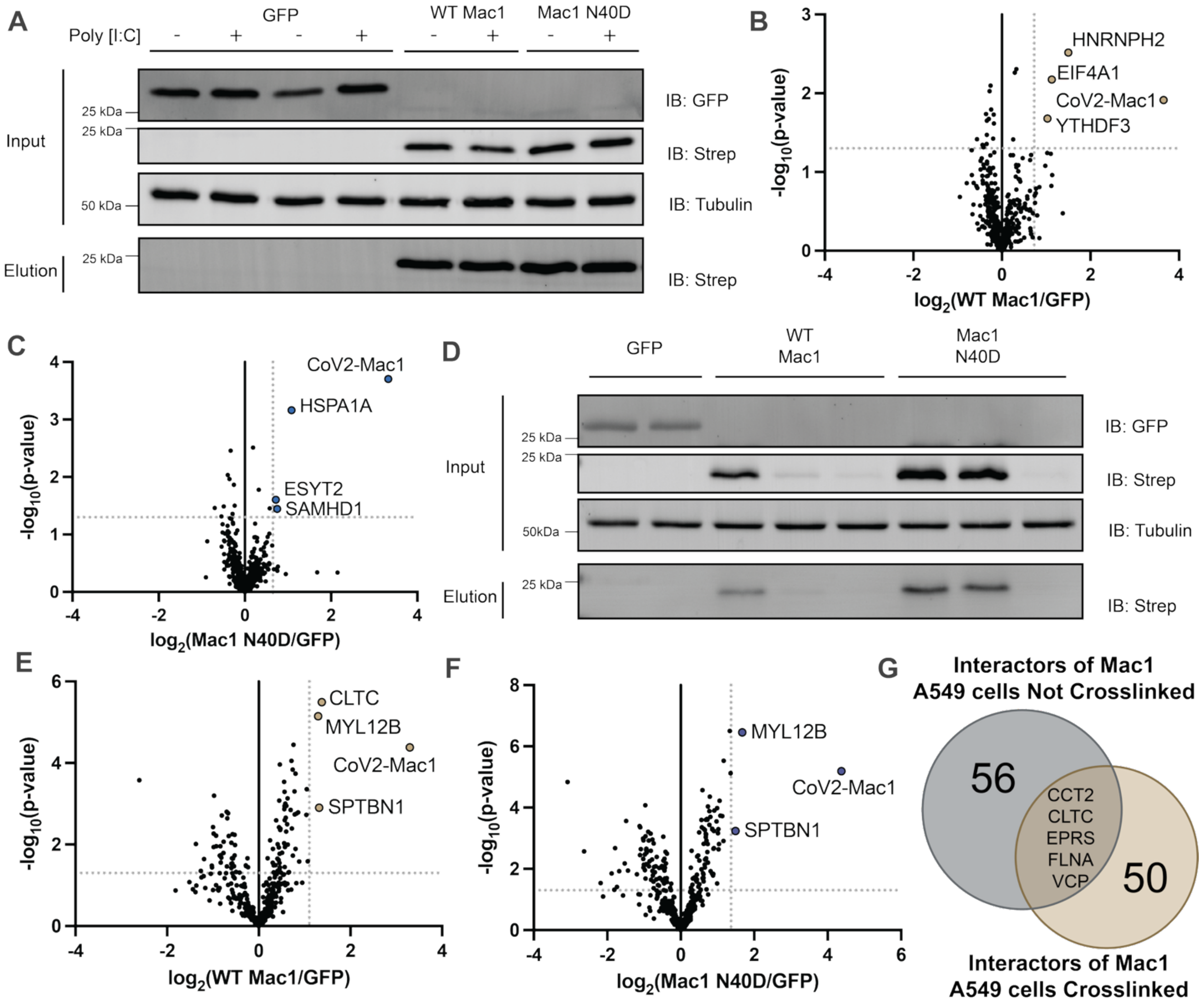


**Figure S3:** Mac1 enrichment from A549 cells. A) Representative Western blot showing signal from GFP, Strep-Tag (representative of Mac1 abundance), and tubulin (loading control) from samples without DSP crosslinking before (input) and after (elution) enrichment for Strep-Tag. A549 cells were transiently transfected with GFP, WT Mac1 or Mac1 N40D. Co-immunoprecipitation was then performed to enrich Strep-Tag and crosslinked interactors. Strep signal in the elution sample in the WT Mac1 and Mac1 N40D lanes indicates that the enrichment was successful. B) Volcano plot comparing identified proteins between WT Mac1 and GFP samples from A549 cells without DSP crosslinking. Cutoffs drawn at 2 SD for enrichment and p-value less than 0.05. Cutoffs of 1SD for enrichment and no p-value restriction were used for analysis. C) Volcano plot comparing identified proteins between Mac1 N40D and GFP samples from A549 cells without DSP crosslinking. Cutoffs drawn at 2 SD for enrichment and p-value less than 0.05. Cutoffs of 1SD for enrichment and no p-value restriction were used for analysis. D) Representative Western blot showing signal from GFP, Strep-Tag (Mac1), and tubulin (loading control) from samples with DSP crosslinking before (input) and after (elution) enrichment for Strep-Tag. A549 cells were transfected and enriched as described in (A). Expression of WT Mac1 and Mac1 N40D was variable between biological replicates. Replicates with comparable high expression (first lane of WT and second and third lanes of N40D in this blot) were taken forward through proteomics analysis. E) Volcano plot comparing identified proteins between WT Mac1 and GFP samples from A549 cells with DSP crosslinking. Cutoffs drawn at 2 SD for enrichment and p-value less that 0.05. Cutoffs of 1SD for enrichment and no p-value restriction were used for analysis. F) Volcano plot comparing identified proteins between Mac1 N40D and GFP samples from A549 cells with DSP crosslinking. Cutoffs drawn at 2 SD for enrichment and p-value less that 0.05. Cutoffs of 1SD for enrichment and no p-value restriction were used for analysis. G) Venn diagram comparing proteins enriched more than 1SD between crosslinked and un-crosslinked samples from A549 cells. CCT2, CLTC, EPRS, FLNA, and VCP were enriched in both crosslinked and un-crosslinked samples.
